# *fastp*: an ultra-fast all-in-one FASTQ preprocessor

**DOI:** 10.1101/274100

**Authors:** Shifu Chen, Yanqing Zhou, Yaru Chen, Jia Gu

**Affiliations:** HaploX Biotechnology; Shenzhen Institutes of Advanced Technology, Chinese Academy of Sciences

## Abstract

**Motivation:** Quality control and preprocessing of FASTQ files are essential to providing clean data for downstream analysis. Traditionally, a different tool is used for each operation, such as quality control, adapter trimming, and quality filtering. These tools are often insufficiently fast as most are developed using high-level programming languages (e.g., Python and Java) and provide limited multi-threading support. Reading and loading data multiple times also renders preprocessing slow and I/O inefficient.

**Results:** We developed *fastp* as an ultra-fast FASTQ preprocessor with useful quality control and data-filtering features. It can perform quality control, adapter trimming, quality filtering, per-read quality cutting, and many other operations with a single scan of the FASTQ data. It also supports unique molecular identifier preprocessing, poly tail trimming, output splitting, and base correction for paired-end data. It can automatically detect adapters for single-end and paired-end FASTQ data. This tool is developed in C++ and has multi-threading support. Based on our evaluation, *fastp* is 2–5 times faster than other FASTQ preprocessing tools such as Trimmomatic or Cutadapt despite performing far more operations than similar tools.

**Availability and Implementation:** The open-source code and corresponding instructions are available at https://github.com/OpenGene/fastp

## 1. Introduction

Quality control and preprocessing of sequencing data are critical to obtaining high-quality and high-confidence variants in downstream data analysis. Data can suffer from adapter contamination, base content biases, and overrepresented sequences. Even worse, library preparation and sequencing steps always involve errors and can cause inaccurate representations of original nucleic acid sequences. Sequencing technologies, especially next-generation sequencing (NGS), have been broadly used in clinical applications in recent years, particularly for noninvasive prenatal testing (Bianchi et al., 2015) and cancer diagnosis. For example, liquid biopsy technology (Esposito, Criscitiello, Trapani, & Curigliano, 2017), which seeks out cancer-related biomarkers in the circulatory system, can be used to facilitate cancer diagnosis and personalized treatment regimens. As a major technology in liquid biopsy, cell-free tumor DNA (ctDNA) sequencing is used to detect tumor-derived DNA fragments from plasma, urine, and other circulating liquids. ctDNA sequencing data are usually highly noisy, and detected mutations often exhibit ultra-low mutation allele frequencies (MAF); quality control and data preprocessing are especially important for detecting low-MAF mutations to eliminate false positives and false negatives.

Quality control and preprocessing of FASTQ data could be considered resolved given the availability of several relevant tools. For instance, FASTQC (Andrews) is a Java-based quality control tool providing per-base and per-read quality profiling features. Cutadapt (Martin, 2011) is a commonly used adapter trimmer, which also provides some read-filtering features. Trimmomatic (Bolger, Lohse, & Usadel, 2014), another popular trimming adapter tool, can perform quality pruning using algorithms such as sliding window cutting. SOAPnuke (Y. Chen et al., 2018) is a recently published tool for adapter trimming and read filtering with the implementation of MapReduce on Hadoop systems.

In the past, multiple tools were employed for FASTQ data quality control and preprocessing. A typical combination was the use of FASTQC for quality control, Cutadapt for adapter trimming, and Trimmomatic for read pruning and filtering. The requirement to read and load data multiple times made preprocessing slow and I/O inefficient. Yet the tools must be used in combination because no single tool currently exists that can effectively address all these problems. The authors developed AfterQC (S. Chen et al., 2017) to integrate quality control, adapter trimming, data filtering, and other useful functions into one tool. AfterQC is a convenient tool that can perform all necessary operations and output HTML-based reports with a single scan of FASTQ files. It also provides a novel algorithm to correct bases by searching for overlapping paired-end reads. However, because AfterQC was developed in Python, it is relatively slow and overly time-consuming when processing large FASTQ files.

In this paper, we present *fastp*, an ultra-fast tool to perform quality control, read filtering, and base correction for FASTQ data. It includes most features of FASTQC + Cutadapt + Trimmomatic + AfterQC while running 2–5 times faster than any of them alone. In addition to the functions available in these tools, *fastp* offers supplementary features such as unique molecular identifier (UMI) preprocessing, per-read polyG tail trimming, and output splitting. *fastp* also provides quality control reports for pre- and post-filtered data within a single HTML page, which allows for direct comparison of quality statistics altered by preprocessing. *fastp* can automatically detect adapter sequences for single-end and paired-end Illumina data. In contrast with the aforementioned tools developed in Java or Python, *fastp* is developed in C/C++ with solid multi-threading implementation, making it much faster than its peers. Furthermore, based on the functions for correcting or eliminating sequencing errors, *fastp* can obtain even better clean data compared to conventional tools.

## 2. Methods

As an all-in-one FASTQ preprocessor, *fastp* provides functions including quality profiling, adapter trimming, read filtering, and base correction. It supports both single-end and paired-end short read data and also provides basic support for long-read data, which are typically generated by PacBio and Nanopore sequencers. In this section, we will first present the overall design of this tool and then explain how the major modules work.

### 2.1 Overall design

*fastp* is designed for multi-threading parallel processing. Reads loaded from FASTQ files will be packed with a size of *N* (*N* = 1000). Each pack will be consumed by one thread in the pool, and each read of the pack will be processed. Each thread has an individual context to store statistical values of the reads it processes, such as per-cycle quality profiles, per-cycle base contents, adapter trimming results, and k-mer counts. These values will be merged after all reads are processed, and a reporter will generate reports in HTML and JSON formats. *fastp* reports statistical values for pre-filtering and post-filtering data to facilitate comparisons of changes in data quality after filtering is complete.

*fastp* supports single-end (SE) and paired-end (PE) data. While most steps of SE and PE data processing are similar, PE data processing requires some additional steps such as overlapping analysis. For the sake of simplicity, we only demonstrate the main workflow of paired-end data preprocessing, shown in Fig. 1.

**Fig. 1.**
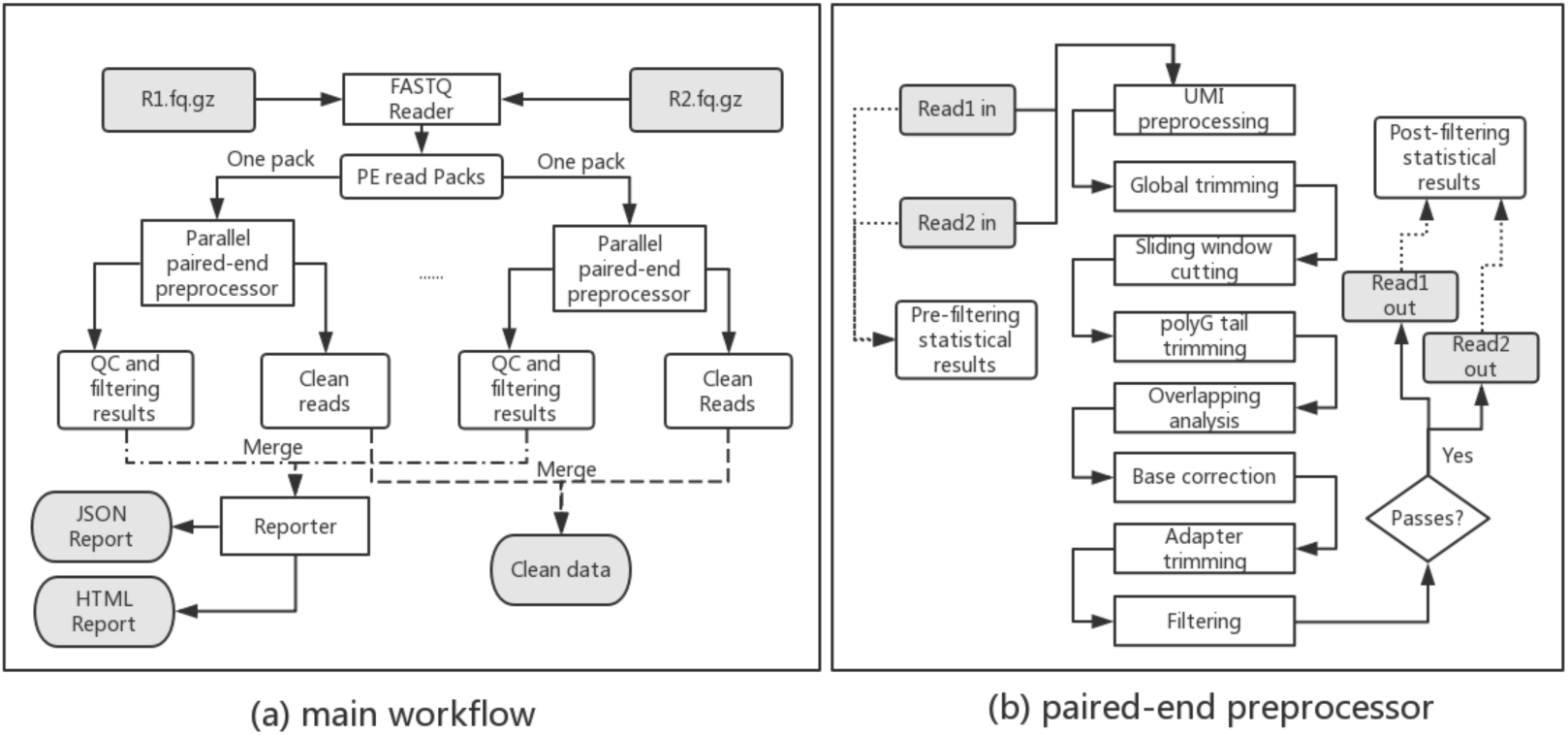
Workflow of *fastp*. (a) Main workflow of paired-end data processing, and (b) paired-end preprocessor of one read pair. In the main workflow, a pair of FASTQ files is loaded and packed, after which each read pair is processed individually in the paired-end preprocessor, described in (b).

### 2.2 Adapter trimming

*fastp* supports automatic adapter trimming for both single-end and paired-end Illumina data and uses different algorithms for each of these tasks. For single-end data, adapter sequences are detected by assembling the high-frequency read tails; for paired-end data, adapter sequences are detected by finding the overlap of each pair.

The adapter-sequence detection algorithm is based on two assumptions: the first is that only one adapter exists in the data; the second is that adapter sequences exist only in the read tails. These two assumptions are valid for major next-generation sequencers like Illumina HiSeq series, NextSeq series, and NovaSeq series. We compute the k-mer (*k* = 10) of first *N* reads (*N* = 1M). From this k-mer, the sequences with high occurrence frequencies (> 0.0001) are considered as adapter seeds. Low-complexity sequences are removed because they are usually caused by sequencing artifacts. The adapter seeds are sorted by its occurrence frequencies. A tree-based algorithm is applied to extend the adapter seeds to find the real complete adapter, which is described by the pseudo code in Algorithm 1.

#### Algorithm 1: adapter sequence detection

~~~
**for** *seed* **in** *sorted_adapter_seeds*:
     *seqs_after_seed* = get_seqs_after(*seed*)
     *forward_tree* = build_nucleotide_tree(*seqs_after_seed*)
     *found* = **True**
     *node = forward_tree*.root
     *after_seed* = “”
     **while** *node*.is_not_leaf():
        **if** *node*.has_dominant_child():
             *node = node*.dominant_child()
             *after_seed = after_seed + node.base*
        **else**:
             *found* = **False**
             **break**
    **if** *found* == **False**:
        **continue**
    **else**:
        *seqs_before_seed* = get_seqs_before(*seed*)
        *backward_tree* = build_nucleotide_tree(*seqs_before_seed*)
        *node = backward_tree*.root
        *before_seed* = “”
        **while** *node*.is_not_leaf():
           **if** *node*.has_dominant_child():
               *node* = node.dominant_child()
               *before_seed* = *node*.base + *before_seed*
           **else**:
               **break**
   *adapter = before_seed + seed + after_seed*
   **break**
~~~

In Algorithm 1, the function build_nucleotide_tree() is used to convert a set of sequences to a tree, in which each node is a nucleotide and each path of root to leaf is a sequence. A node’s dominant child is defined as its major child with a dominant percentage (>90%). This algorithm tries to extend an adapter seed in the forward direction to check its validity since a valid adapter can always be extended to the read tails. And if this adapter seed is valid, a backward extension is applied to obtain the complete adapter sequence. The process of extending an adapter seed in forward and backward directions is given in Fig. 2.

**Fig. 2.**
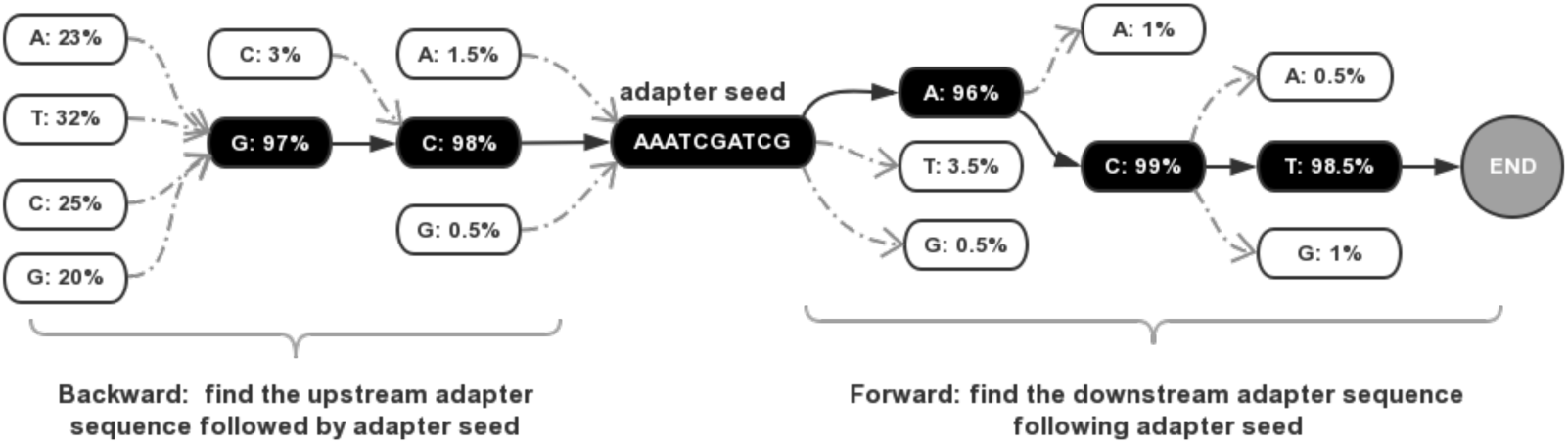
A demonstration of extending an adapter seed in both forward and backward directions. The found adapter is GCAAATCGATCGACT, with the first two bases (GC) as the upstream sequence, the central ten bases as the adapter seed, and the last three bases (ACT) as the downstream sequence.

For paired-end data, *fastp* seeks the overlap of each pair and considers the bases that fall out of the overlapped regions as adapter contents. The overlapping detection algorithm was derived from our previous work, AfterQC. Compared to sequence-matching-based adapter-trimming tools like Cutadapt and Trimmomatic, a clear advantage of the overlap-analysis-based method is that it can trim adapters with few bases in the read tail. For example, most sequence-matching-based tools require a hatchment of at least three bases and cannot trim adapters with only one or two bases. In contrast, *fastp* can trim adapters with even only one base in the tail.

Although *fastp* can detect adapter sequences automatically, it also provides interfaces to set specific adapter sequences for trimming. For SE data, if an adapter sequence is given, then automatic adapter-sequence detection will be disabled. For PE data, the adapter sequence will be used for sequence-matching-based adapter trimming only when *fastp* fails to detect a good overlap in the pair.

### 2.3 Base correction

For paired-end data, if one pair of reads can be detected with a good overlap, then the bases within the overlapped region can be compared. If the reads are of high quality, they are usually completely reverse-complemented.

If any mismatches are found within the overlapped region, *fastp* will try to correct them. *fastp* only corrects a mismatched base pair with an imbalanced quality score, such as when one has a high-quality score (> Q30) and the other has a low-quality score (< Q15). To reduce false corrections, *fastp* only performs a correction if the total mismatch is below a given threshold *T* (*T* = 5).

### 2.4 Sliding window quality cutting

To improve the read quality, *fastp* supports a sliding window method to drop the low-quality bases of each read’s head and tail. The window can slide from either 5’ to 3’ or from 3’ to 5’, and the average quality score within the window is evaluated. If the average quality is lower than a given threshold, then the bases in the window will be marked as discarded and the window will be moved forward by one base; otherwise, the algorithm ends.

### 2.5 polyG and polyX tail trimming

PolyG is a common issue observed in Illumina NextSeq and NovaSeq series, which are based on two-color chemistry. Such systems use two different lights (i.e., red and green) to represent four bases: a base with only a detected red-light signal is called C; a base with only a detected green light signal is called T; a base with both red and green light detected is called A; and a base with no light detected is called G. However, as the sequencing by synthesis proceeds to subsequent cycles, the signal strength of each DNA cluster becomes progressively weaker. This issue causes some T and C to be wrongly interpreted as G in the read tails, a problem otherwise known as a polyG tail.

*fastp* can detect and trim the polyG in the read tails. It checks the flow cell identifier to determine whether the data are from Illumina NextSeq or NovaSeq sequencers, and if so, it automatically enables polyG tail trimming. The polyG tail issue can result in a serious base content separation problem, meaning that A and T or C and G have substantially different base content ratios. Fig. 3 shows an example of data exhibiting a polyG tail issue and how the problem is addressed with *fastp* preprocessing.

**Fig. 3.**
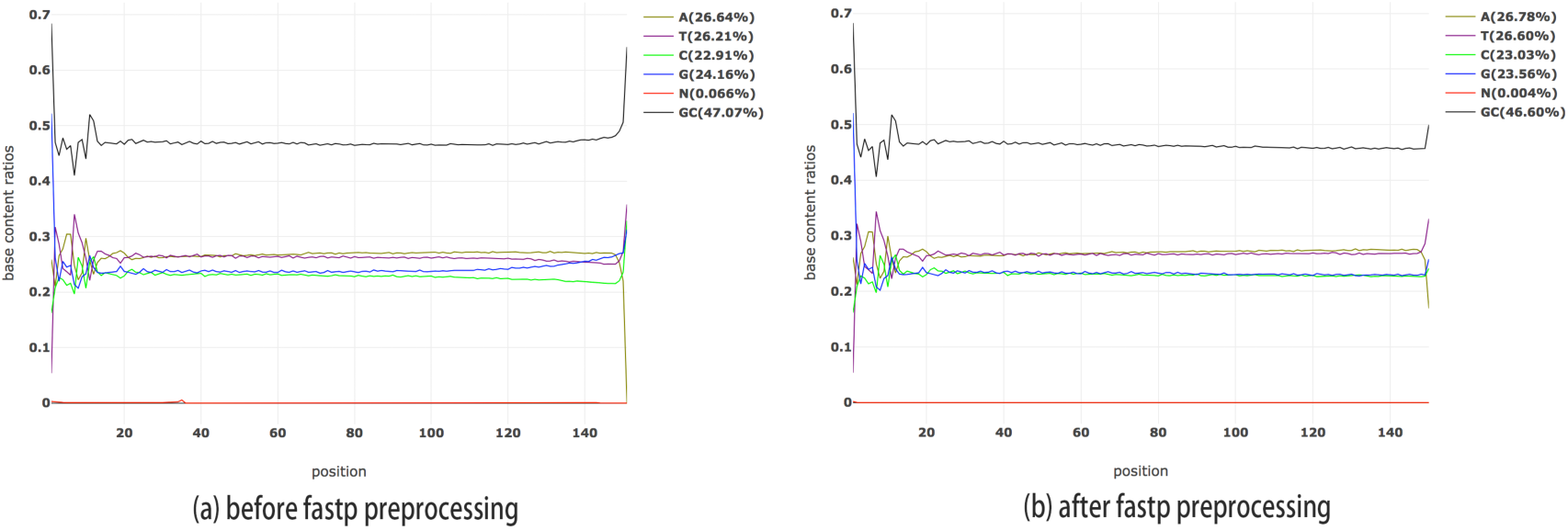
The base content ratio curves generated by *fastp* for one Illumina NextSeq FASTQ file. (a) before *fastp* preprocessing, and (b) after *fastp* preprocessing. As depicted in (a), the G curve is abnormal, and the G/C curves are separated. In (b), the G/C separation problem is eliminated.

*fastp* also implements polyX tail trimming, where X means any base of A/T/C/G. This function can be used to trim the low-- complexity consecutive bases in 3’ end. PolyX tail trimming and polyG tail trimming can be enabled together. In this case, polyG tail trimming will be applied first since polyG is usually caused by sequencing artifacts and should be trimmed first.

### 2.6 UMI preprocessing

Recently, UMI technology was proposed to reduce background noise and improve sensitivity when detecting ultra-low frequency mutations in deep-sequencing applications (i.e., ctDNA sequencing). The UMI method can be used to remove duplications and generate high-quality consensus reads. It has been adopted by various sequencing methods such as Duplex-Seq (Kennedy et al., 2014) and iDES (Newman et al., 2016). For Illumina sequencing platforms, UMI can be integrated into the sample index or inserted DNA. UMI should be shifted to the read identifier to be retained by alignment tools like BWA (Li & Durbin, 2009) or Bowtie (Langmead & Salzberg, 2012).

Some tools have already been developed for preprocessing UMI-integrated FASTQ data, such as UMI-tools (Tom Sean Smith, 2018) and umis (Valentine Svensson, 2018). However, these tools are not efficient enough and require individual execution that consumes additional I/O and computational resources. *fastp* supports UMI preprocessing with little overhead; that is, it supports UMI in either a sample index or inserted DNA (or both). Compared to UMI-tools or umis, *fastp* runs approximately 3 times faster even when performing other tasks simultaneously (i.e., QC and filtering). Performance evaluation results are discussed in the next section.

### 2.7 Output splitting

Parallel processing of NGS data has become a new trend, especially in a cloud-computing environment. In a typical parallel NGS data processing pipeline, an original FASTQ file will be split into multiple pieces, and each piece will be run with aligners and alignment adjustment tools to obtain the corresponding BAM file. These BAM files can then be merged into different forms for parallel variant calling.

*fastp* supports two splitting modes: splitting by file lines and splitting by file numbers. The latter is more complicated because *fastp* must evaluate the total lines of the input files, which is especially difficult for GZIP-compressed data. *fastp* evaluates total lines by comparing the stream size of the first 1M reads.

### 2.8 Overrepresented sequence analysis

Some sequences, or even entire reads, can be overrepresented in FASTQ data. Analysis of these overrepresented sequences provides an overview of certain sequencing artifacts such as PCR over-duplication, polyG tails, and adapter contamination. FASTQC offers an overrepresented sequence analysis module; however, according to the author’s introduction, FASTQC only tracks the first 1M reads of the input file to conserve memory. We suggest that inferring the overall distribution from the first 1M reads is not a reliable solution as the initial reads in Illumina FASTQ data usually originate from the edges of flowcell lanes, which may have lower quality and different patterns than the overall distribution.

Unlike FASTQC, *fastp* samples all reads evenly to evaluate overrepresented sequences and eliminate partial distribution bias. To achieve efficient implementation of this feature, we designed a two-step method. In the first step, *fastp* completely analyzes the first 1.5M base pairs of the input FASTQ to obtain a list of sequences with relatively high occurrence frequency in different sizes. In the second step, *fastp* samples the entire file and counts the occurrence of each sequence. Finally, the sequences with high occurrence frequency are reported.

Besides the occurrence frequency, *fastp* also records the positions of overrepresented sequences. This information is quite useful for diagnosing sequence quality issues. Some sequences tend to appear in the read head whereas others appear more often in the read tail. The distribution of overrepresented sequences is visualized in the HTML report. Fig. 4 shows a demonstration of overrepresented sequence analysis results.

**Fig. 4.**
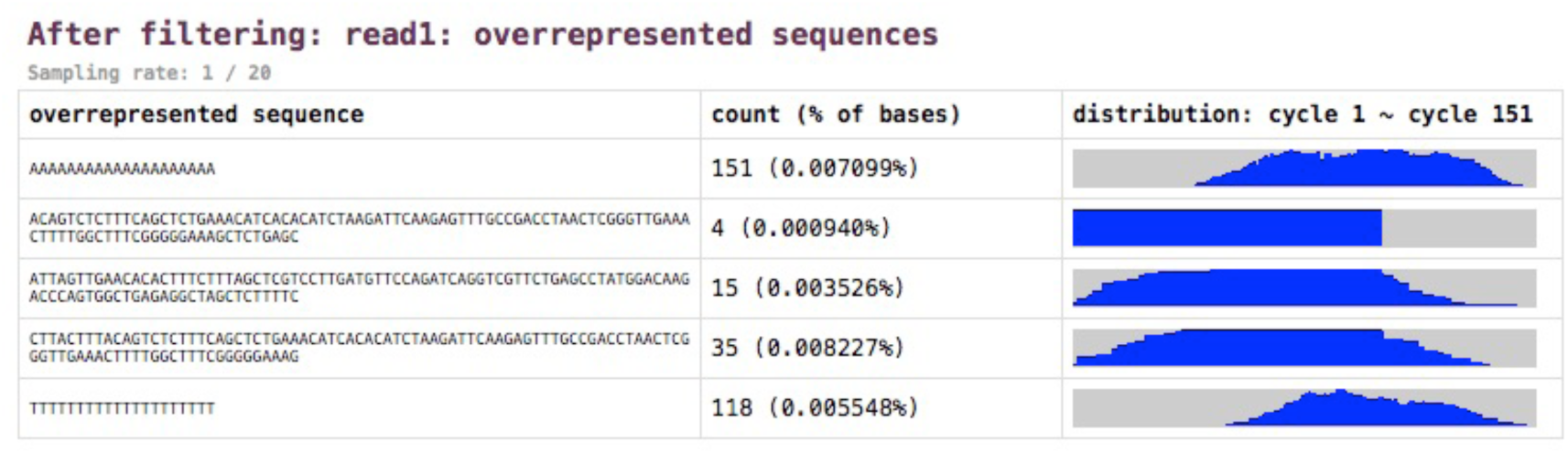
Overrepresented sequence analysis results. The right column shows the histogram of occurrence among all sequencing cycles.

### 2.9 Quality control and reporting

*fastp* supports filtering reads using a low-quality base percentage, *N* base number, and read length. These filters are trivial and thus not described here. *fastp* also supports filtering low-complexity reads by evaluating the percentage of consecutive bases. *fastp* records the number of reads that were filtered out according to different filtering criteria.

*fastp* also provides comprehensive information on quality-profiling results. In contrast to FASTQC, *fastp* offers results for both pre-filtering and post-filtering data, which allows for evaluation of the filtering effect by comparing the figures directly. *fastp* also reports results in JSON and HTML format, the former of which contains all data visualized in the HTML report. The format of the JSON report is manually optimized to be easily readable by humans. The HTML report is a single standalone web page, with all figures created dynamically using JavaScript and web canvas. Additionally, *fastp* provides a full k-mer occurrence table for all 6-bp sequences. An online demonstration of the HTML report can be found at http://opengene.org/fastp/fastp.html.

## 3. Results

We conducted several experiments to evaluate the performance of *fastp* in terms of speed and quality. We chose FASTQC, Cutadapt, Trimmomatic, SOAPnuke, and AfterQC for performance comparison, and the results revealed that *fastp* is much faster than these tools while providing similar or even better quality.

### 3.1 Speed evaluation

We compared the speed of all six tools by preprocessing the B17NCB1 dataset, obtained from the National Center for Clinical Laboratories in China. This dataset is paired-end, containing 9,316 M bases. We evaluated the used time for PE and SE mode, respectively. All tools were run with a single thread to ensure fair comparison. Results are listed in Table 1.

**Table 1.** Speed comparison of *fastp* and other software.

| Tools | PE / SE | Time (min) | Throughput (read/s) |
| --- | --- | --- | --- |
| fastp | SE | 6 | 129397.9 |
|  | PE | 13.3 | 116750.0 |
| FASTQC | SE | 13 | 59722.1 |
|  | PE | 25.8 | 60185.1 |
| Cutadapt | SE | 18 | 43132.6 |
|  | PE | 24.6 | 63120.9 |
| SOAPnuke | SE | 30.7 | 25289.5 |
|  | PE | 32.5 | 47777.7 |
| AfterQC | SE | 25.2 | 30809.0 |
|  | PE | 57.2 | 27146.4 |
| Trimmomatic | SE | 31.9 | 24338.2 |
|  | PE | 60.9 | 25497.1 |

As indicated, *fastp* is much faster than other tools. The second-fastest tool is FASTQC, which takes about twice the time of *fastp*. However, FASTQC only performs quality control, whereas *fastp* performs quality control (for pre-filtering data and post-filtering data), data filtering, and other operations. The other tools take 3–5 times longer than *fastp*. Because *fastp* was natively designed for multi-thread processing, it may demonstrate even higher performance when executed in real applications in multi-thread mode. As some of the selected tools do not support multi-threading, a multi-threading performance comparison is not provided here.

### 3.2 Quality evaluation

To evaluate the adapter trimming and quality pruning of *fastp* compared to other tools (i.e., AfterQC, SOAPnuke, Trimmomatic, and Cutadapt), we used an Illumina NextSeq PE150 dataset (NS_PE150). Among these tools, *fastp* and AfterQC can trim adapters via overlap analysis, whereas the other tools require adapter sequence input. We evaluated the amount of suspected adapters by searching the 33bp-long adapter sequences from post-filtering data with tolerance of several mismatches; results are shown in Fig. 5.

**Fig. 5.**
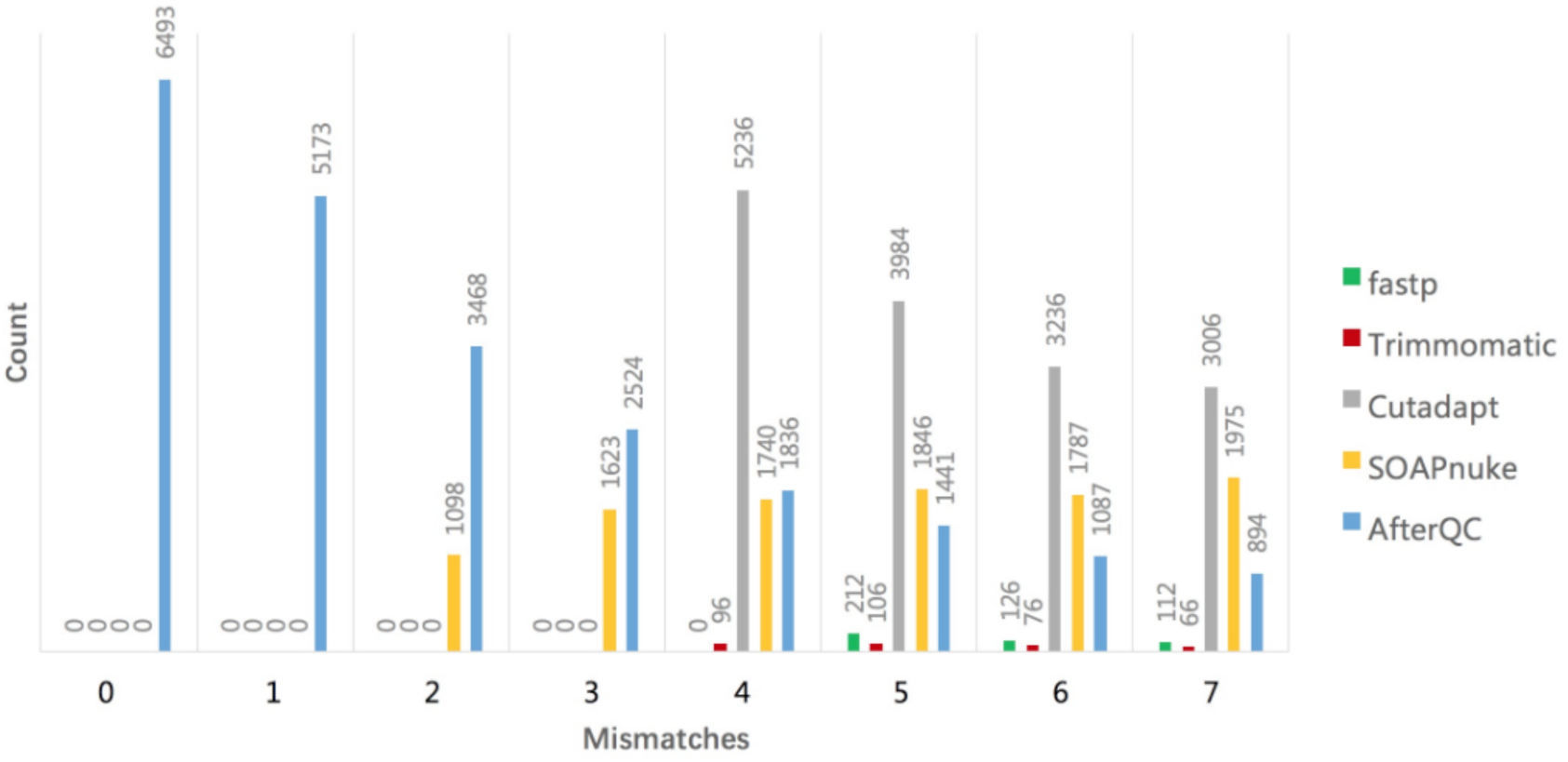
Result of adapter trimming performance evaluation. The X-axis is the number of allowed mismatches when searching for suspected adapter sequences, and the Y-axis is the count of suspected adapter sequences.

From Fig. 5, we can learn that *fastp* and Trimmomatic give the best performance. The *fastp*-filtered data contains no suspected adapters when 4 or fewer mismatches are allowed. Comparing to *fastp*-filtered data, Trimmomatic-filtered data contains less suspected adapters when 5 or more mismatches are allowed, but contains more when 4 mismatches are allowed. The data filtered with Cutadapt, SOAPnuke or AfterQC contain a large number of suspected adapters when the allowed mismatches are 4 or more. Since AfterQC was not designed as a professional adapter-trimming tool, the data filtered by this tool even contains suspected adapters without any mismatch.

To further evaluate filtering effectiveness, we mapped the data filtered by different tools to the reference genome hg19 using BWA-MEM and evaluated the mapping results with Samtools (Li et al., 2009). Mismatches, clips, and improper mappings were recorded for evaluation purposes. From our perspective, uncorrected sequencing errors contribute greatly to mismatches while residual adapters contribute to clips and improper mappings. Comparison results are presented in Table 2.

**Table 2.** Mismatches, clips, and single-read maps of the data filtered with different tools.

| NS_PE150 | Total map<br>base (M) | Mismatched<br>base (M) | Total map read<br>(M) | Clip read (M) | Single-read map |
| --- | --- | --- | --- | --- | --- |
| Raw data | 3390 | 19.8 | 22.4 | 6.16 | 46025 |
| *fastp* | 3102 | 10.6 | 21.2 | 0.30 | 271 |
| AfterQC | 3053 | 12.3 | 21.2 | 0.64 | 34099 |
| SOAPnuke | 2981 | 18.3 | 19.7 | 3.46 | 36888 |
| Trimmomatic | 3111 | 12.8 | 22.1 | 0.75 | 229111 |
| Cutadapt | 3287 | 19.8 | 22.4 | 1.05 | 46195 |

*fastp* generated the lowest number of mismatches, clipped reads, and single-read mapped reads. Trimmomatic and Cutadapt generated much more clipped or single-read mapped reads. Given that Trimmomatic, Cutadapt, and SOAPnuke are all based on adapter-sequence matching, they may fail to detect adapters when the adapter sequence only has a few bases. For example, Cutadapt requires at least 3bp matching of the adapter sequence and the read for a sequence to be recognized as an adapter. If the adapter sequence has only one or two bases, it will not be detected and is often erroneously reported as either a mismatch or soft clip by the alignment tools.

### 3.3 UMI evaluation

UMI technology is widely used in cancer sequencing, especially ctDNA sequencing. To analyze NGS data with UMI integration, the FASTQ preprocessor should shift the UMI from the reads to the read identifiers. We ran UMI preprocessing on a FASTQ of 4Gb Illumina PE150 data using *fastp*, umis, and UMI-tools, respectively. The execution times were recorded and are reported in Table 3.

**Table 3.**
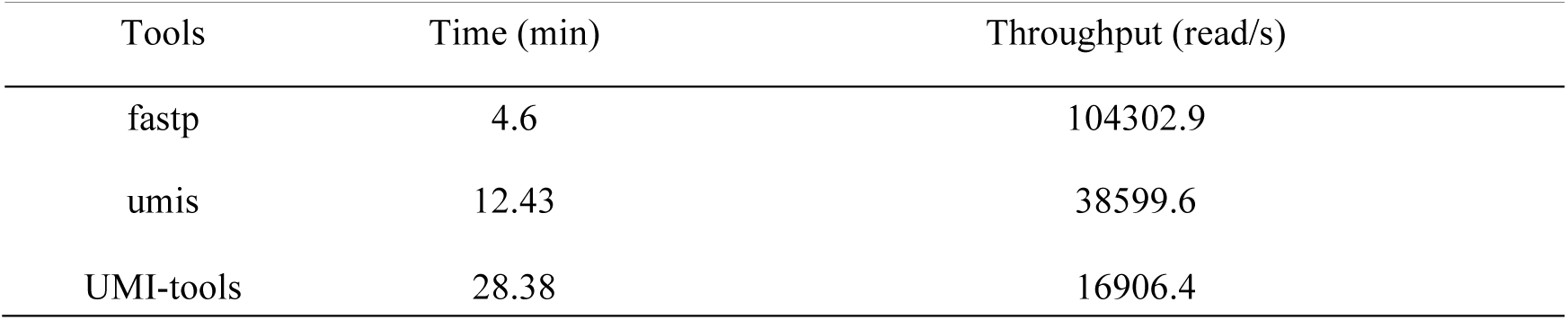
UMI preprocessing time comparison of *fastp*, umis, and UMI-tools.

Clearly, *fastp* is approximately 2.7 times faster than umis and about 6.1 times faster than UMI-tools. This evaluation was conducted with GZIP input and uncompressed output because umis does not support GZIP output. Considering that *fastp* can achieve high performance for UMI preprocessing, it has recently been adopted by the popular NGS pipeline framework, bcbio-nextgen (Brad Chapman, 2018).

## 4. Discussion

In this paper, we introduced *fastp*, an ultra-fast all-in-one FASTQ preprocessor. *fastp* is a versatile tool that can perform quality profiling, read filtering, read pruning, adapter trimming, polyG/polyX tail trimming, UMI preprocessing, and other operations with a single scan of FASTQ files. Additionally, it can split output into multiple files for parallel processing.

We evaluated the performance of speed and quality of *fastp* against other tools. The results indicate that *fastp* is much faster than its counterparts and provides the highest-quality data filtering of all other tested options. *fastp* is an open-source software. Due to its high speed and excellent quality in FASTQ file quality control and filtering, *fastp* has gained many community users.

## 5. Acknowledgement

The authors would like to thank *fastp* community users for identifying bugs.

